# Assessing and enhancing foldability in designed proteins

**DOI:** 10.1101/2021.11.09.467863

**Authors:** Dina Listov, Rosalie Lipsh-Sokolik, Stéphane Rosset, Che Yang, Bruno E Correia, Sarel J Fleishman

## Abstract

Recent advances in protein-design methodology have led to a dramatic increase in reliability and scale. With these advances, dozens and even thousands of designed proteins are automatically generated and screened. Nevertheless, the success rate, particularly in design of functional proteins, is low and fundamental goals such as reliable *de novo* design of efficient enzymes remain beyond reach. Experimental analyses have consistently indicated that a major reason for design failure is inaccuracy and misfolding relative to the design model. To address this challenge, we describe complementary methods to diagnose and ameliorate suboptimal regions in designed proteins: first, we develop a Rosetta atomistic computational mutation scanning approach to detect energetically suboptimal positions in designs (available on a web server https://pSUFER.weizmann.ac.il); second, we demonstrate that AlphaFold2 *ab initio* structure prediction flags regions that may misfold in designed enzymes and binders; and third, we focus FuncLib design calculations on suboptimal positions in a previously designed low-efficiency enzyme improving its catalytic efficiency by 330 fold. Furthermore, applied to a *de novo* designed protein that exhibited limited stability, the same approach markedly improved stability and expressibility. Thus, foldability analysis and enhancement may dramatically increase the success rate in design of functional proteins.

## Introduction

Over the past decade, protein design methodology has made remarkable progress. New methods enable the routine design of new folds ^1–4^, assemblies ^5,6^, and new or improved functions ^7–13^. Despite these achievements, however, only a small fraction of experimentally tested designs are functional ^8,14,15^, and fundamental protein design objectives still lie beyond our reach. For example, reliable *de novo* enzyme design remains an unsolved challenge despite decades of research, and to date, all designs exhibit low efficiency ^15–17^. Previous analyses of successful and failed enzyme designs indicated that inaccuracies in the design of catalytic constellations are partly to blame for failures ^18–21^. Furthermore, observations across many different fold, binder, and enzyme design studies indicated misfolding relative to the design model as likely to be the most general and critical problem in failed designs ^2,22–25^. Indeed, the significance of misfolding has served as the primary motivation for studies regarding the principles for designing new and idealized folds ^2,26,27^ and for developing general methods to improve designed proteins’ foldability and stability ^11,13,27–29^.

The current study is motivated by our ongoing efforts to develop a reliable strategy for *de novo* enzyme design. Our strategy is based on recent methods developed in our lab to design new backbones through the modular assembly of large (60-150 amino acid) backbone fragments of natural enzymes ^25,30,31^. Modular assembly and design was successful in generating accurate antibodies ^27^ and ultrahigh specificity binders ^24^ and enzymes ^25^ with as many as 100 mutations from any natural protein. Applied to *de novo* enzyme design, however, this same strategy has so far failed to generate high-efficiency enzymes. Furthermore, the majority of the failed designs exhibited non-cooperative folding transitions. The correspondence between foldability and activity in our ongoing *de novo* enzyme design study prompts us to examine the sources of low foldability in proteins generated using current atomistic design methods and to develop strategies for detecting and ameliorating suboptimality in designed proteins.

## Results

### Energy-based suboptimality detection

We develop an automated computational approach to identify energetically suboptimal positions in protein designs based on their model structure. Our approach, which we call pSUFER for protein Strain, Unsatisfactoriness, and Frustration findER, starts by relaxing the input structure using Rosetta atomistic modeling. It then models all single-point mutations at every position, iterating sidechain packing and whole-protein minimization and computes the change in system energy (ΔΔ*G*) relative to the parental protein. These calculations use the Rosetta all-atom energy function 2015 (ref2015) which is dominated by van der Waals packing, hydrogen bonding, electrostatics and implicit solvation ^32^. Finally, pSUFER flags positions that exhibit more than a certain number of mutations (typically 5) that lower the native-state energy (ΔΔ*G* < 0) as potentially suboptimal. These thresholds were chosen empirically and may be changed according to modeling needs. An online web server for automatically running pSUFER calculations is available for academic users at https://pSUFER.weizmann.ac.il. Customizable scripts for automatically relaxing a protein structure, computing suboptimal positions and visualizing them in PyMOL are available in https://github.com/Fleishman-Lab/pSUFER.

Our approach is similar in principle to strategies for computing local frustration in proteins ^33,34^. Methods for analyzing local frustration detect contacting pairs of amino acids that exhibit high energies relative to a computed ensemble of mutational or structural decoys. Frustrated positions are often associated with the protein’s activity since active-site positions and regions that are involved in allosteric communication or conformational change are evolutionarily selected for their role in activity rather than for improving native-state stability ^35^. We note, however, that since these previous approaches search for high-energy pairs of positions, they do not directly detect positions that may be optimized through single-point mutations. By contrast, pSUFER indicates specific positions for design.

### Orders of magnitude improvement in catalytic efficiency of a failed design

To test pSUFER, we analyzed a set of enzymes and binders designed and experimentally characterized over the past four years. To be included in this set, we required that the same design method produced both proteins that accurately folded into the design conception according to X-ray crystallography and ones that misfolded. Despite the availability of experimental structures in these cases, we applied pSUFER to the models to verify that the method could uncover flaws in the design conception without recourse to experimental data. The designs included enzymes ^25^ and binders ^24^ generated through modular backbone assembly and design as well as binders in which an immunogenic epitope was incorporated in a *de novo* designed scaffold protein ^36^. Therefore, these designs encompass a range of contemporary applications of protein design methodology.

The first design pair we examine was generated by modular assembly and design of glycoside hydrolase 10 (GH10) xylanases ^25^. In this design set, 21 out of 43 designs exhibited detectable xylanase activity, and only two exhibited high activity levels as observed in natural enzymes from the GH10 family. From that study, we chose for pSUFER analysis two designs that exhibited the highest and lowest levels of detectable activity and for which we have crystallographic data, xyl3.1 and xyl8.3 (*k*_cat_/K_M_ 9,417 and 0.61 M^-1^s^-1^ respectively). Both designs were highly mutated relative to natural GH10 enzymes, exhibiting 105 and 130 mutations from any natural enzyme, respectively (out of approximately 350 amino acids) and exhibited apparent thermal denaturation temperatures > 55 °C. The crystal structure of xyl3.1 showed remarkable accuracy relative to the design conception with 0.7 Å root mean square deviation (rmsd) across the entire protein and <1 Å all-atom rmsd of active-site residues. By contrast, although xyl8.3 was globally accurate (rmsd = 0.9 Å) and core catalytic groups aligned well between the design model and the experimental structure, the experimental structure exhibited significant missing density in two neighboring loops near the active-site pocket, indicative of local misfolding. Visual inspection of the two designs following their structure determination failed to suggest significant flaws in xyl8.3 that might explain design inaccuracy.

Applied to the two xylanase design models, pSUFER flags a similar number of positions. As expected, most of the flagged positions are either in the active-site pocket or in solvent-accessible positions (Fig 1A,1B). Conspicuously, however, in xyl8.3, pSUFER flags position Lys306, which is buried at the stem of one of the loops that does not exhibit electron density. Furthermore, Lys306 is not stabilized by counter charges, suggesting that the loop disorder may be the result of strain in and around this position.

**Figure 1.**
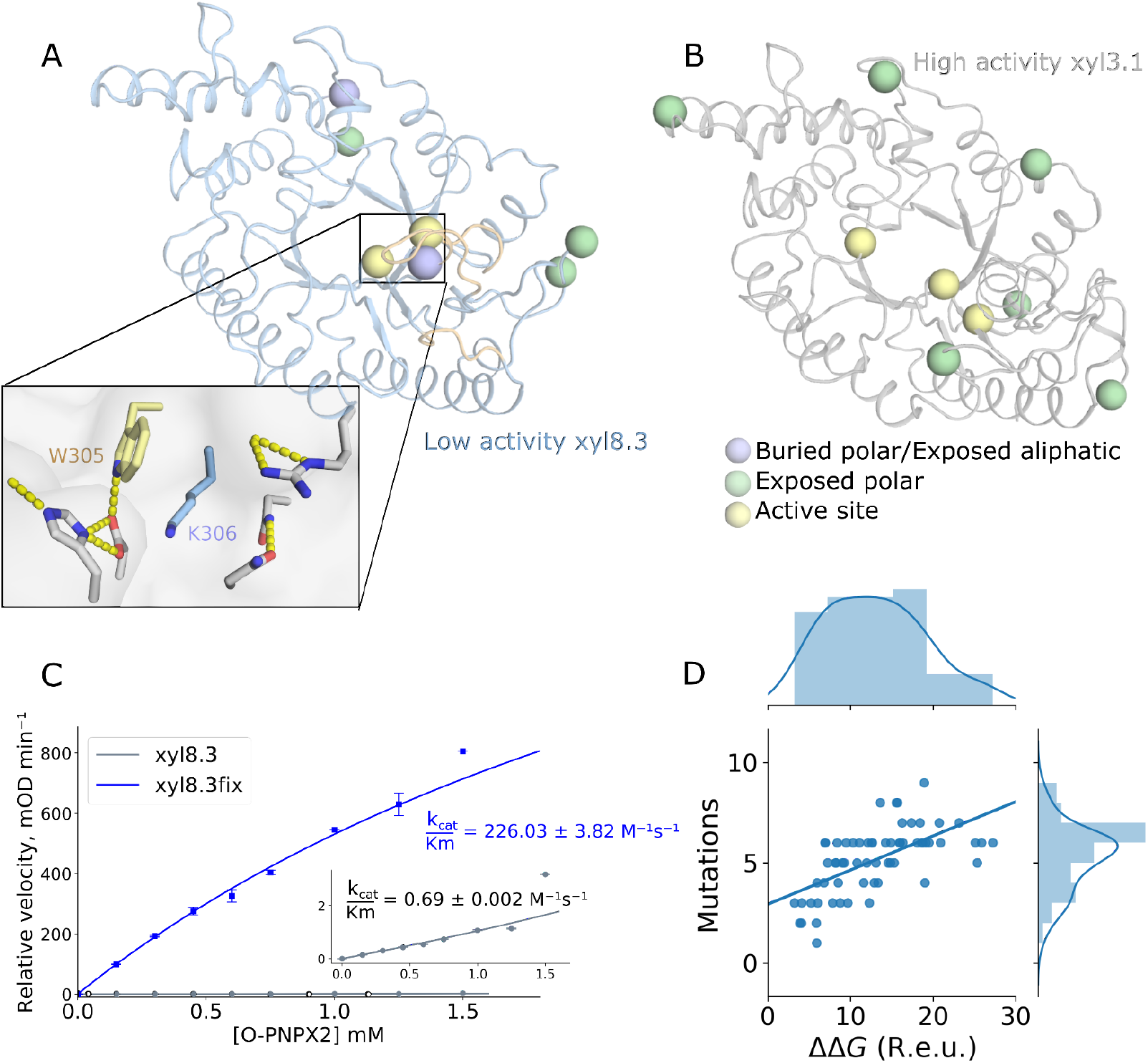
pSUFER analysis of two GH10 xylanases generated using modular assembly and design. **(A)** xyl8.3 backbone shown in cartoon with regions that failed to exhibit electron density in a crystallographic analysis (PDB entry: 6FHE) shown in wheat. (inset) Lys306 is flagged by pSUFER since the Lys is buried in a hydrophobic region without countercharge stabilization. Lys306 is located directly under the two loops that failed to exhibit electron density and is in close contact with active-site position Trp305. **(B)** For comparison, in the case of the high-efficiency and accurately designed xyl3.1 pSUFER only flags surface-exposed polar positions and active-site positions. In both cases, a position was flagged if computational mutation scanning suggested at least six amino acid identities with ΔΔG < 0 at the position. **(C)** xyl8.3fix shows an improvement of 330 fold in activity compared to xyl8.3, *k*_*cat*_/*K*_M_=226 and 0.69 M^-1^s^-1^, respectively. Data points and standard deviations are based on at least two repetitions. **(D)** Improvement in system energy following FuncLib design of positions flagged by pSUFER in 62 models of *de novo* designed enzymes generated by modular assembly and design. On average, 5-6 mutations are introduced, yielding an average improvement of 13 Rosetta energy units (R.e.u.).

We hypothesized that eliminating the strain observed around Lys306 (and the other positions flagged by pSUFER) might improve the enzyme’s catalytic efficiency. To test this hypothesis, we applied FuncLib design ^12^ to the five positions pSUFER flagged outside the active-site pocket in xyl8.3. FuncLib starts by using phylogenetic information and atomistic design calculations to rule out mutations that may be destabilizing to the native-state structure. It then enumerates all combinations of allowed mutations at the designed positions, ranks them by energy and suggests low-energy combinations of mutations. An advantage of the FuncLib methodology over typical stochastic combinatorial design algorithms is that FuncLib relaxes each combination of mutations by whole-protein minimization. Therefore, FuncLib may find stabilizing mutations, including radical small-to-large mutations in the core of the protein that may elude other atomistic design methods ^37–39^. Applied to the five suboptimal positions flagged by pSUFER, one of the top FuncLib designs, which we called xyl8.3fix, improved system energy by 10 Rosetta energy units (R.e.u) by mutating four positions (Thr1Asn, Asn4Glu, Gly139Asp, and Lys306Leu). Among these four, only the Lys306Leu mutation is radical, whereas the others impact solvent-exposed positions to increase surface polarity.

We expressed xyl8.3 and xyl8.3fix fused N-terminally to maltose-binding protein (MBP) in *E. coli* BL21 cells and purified the proteins using an amylose column. We then tested the two designs’ catalytic efficiency using the 4-nitrophenyl β-xylobioside (OPNPX2) chromogenic substrate. Remarkably, Michaelis-Menten analysis revealed that xyl8.3fix exhibits catalytic efficiency of 226 M^-1^s^-1^ (Fig 1C), 330-fold greater than that of the parental xyl8.3. We also analyzed the thermal denaturation of the two designs, finding that both had similar apparent denaturation temperatures 57 and 59 °C for xyl8.3 and xyl8.3fix, respectively (Fig S1). Thus, the dramatic improvement in catalytic efficiency was not due to protein stabilization. Rather, it was likely due either to an improvement in the design’s ability to fold into the active conformation or to reduced strain in the active site.

Significantly, in the modular assembly and design study that included xyl3.1 and xyl8.3 ^25^, we noted that assembling backbone fragments from more than a few natural templates led to low activity (xyl3.1 and 8.3 were based on 3 and 8 backbone fragments, respectively). The activity we observe for xyl8.3fix, however, would place it among the top three designs in that set despite comprising fragments from eight proteins. Thus, the pSUFER analysis may significantly improve the reliability of protein design methods and their ability to generate diverse and functional proteins.

To test whether a similar approach may improve computed system energies in other designs, we applied FuncLib to positions flagged by pSUFER in 62 proteins designed in our lab to generate *de novo* enzymes. FuncLib introduced an average of five mutations and improved the energies by an average of 13 R.e.u in this set (Fig 1D), similar to the values we obtained by applying this approach to xyl8.3.

### Suboptimality in designed binders

We next ask whether pSUFER may shed light on problems in designed binders. In the following example, we examine *de novo* designed binders of biosensors that detect and quantify epitope-specific antibodies. The design strategy uses a bottom-up approach: first defining the function-rendering motifs and then designing a protein fold to support the motifs ^36^. One of the designs (4H.01) forms the binding site accurately as determined by X-ray crystallography; however, this design shows substantial backbone deviations (2.9 Å) relative to the design and exhibits missing density in a loop (Fig 2A). pSUFER flags eleven positions in this design; for instance, Gln84, which is positioned in a cavity next to the loop that exhibits missing density, and Leu78. According to the design model, Gln84 is partly desolvated and does not form intimate polar contacts with the loop. Leu78 is forced into a strained sidechain conformation exhibiting Rosetta sidechain conformation energy (fa_dun) of approximately +3 energy units. This sidechain cannot pack into a favorable conformation due to steric overlaps in relaxed sidechain conformations. In contrast to this design, in the accurately crystallized design, only three positions are flagged, all of which are solvent-exposed (His1, Thr19, and Thr35, Fig 2B).

**Figure 2.**
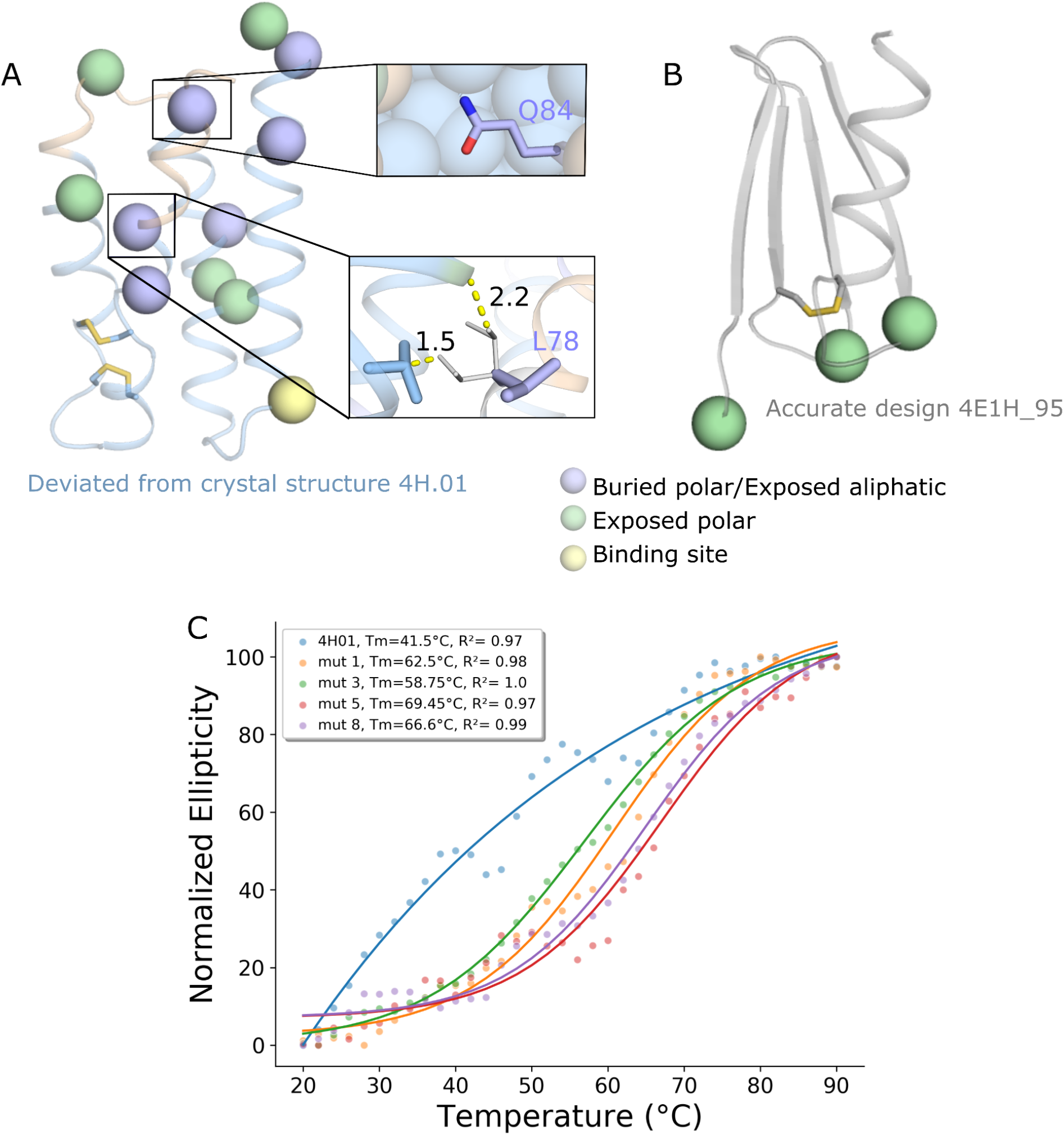
pSUFER analysis of *de novo* designed binders. **(A+B)** pSUFER analysis of *de novo* designed binders. Positions were flagged if mutational scanning suggested at least four amino acid identities with ΔΔ*G* < 0. Wheat backbone marks missing density on the design model. **(A)** Crystallographic analysis of 4H.01 (PDB entry: 6YWD) revealed regions of missing density. pSUFER flags several strained positions surrounding the missing density. (insets) Gln53 is partly desolvated but does not form stabilizing hydrogen bonds. The sidechain conformation of Leu78 is strained in the design model (purple). The most likely sidechain conformation for this position (white) is disallowed due to steric overlaps with neighboring sidechains. **(B)** Design 4E1H_95 was atomically accurate as verified by x-ray crystallography (PDB entry: 6YWC) and pSUFER analysis flags only a few exposed polar sidechains. **(C)** Temperature melts of 4H01 variants monitored by CD at 220 nm. The original 4H.01 does not show cooperative melting curves as opposed to the FuncLib designs.

We applied FuncLib to alleviate strain in the 4H.01 design. We chose eight flagged positions that were far from the binding site (His1, Met2, Glu7, His47, Gln53, Gly76, Leu78, Gln84) for FuncLib design. As 4H.01 is a *de novo* design, phylogenetic analysis, which is a critical part of the FuncLib design strategy ^12^, cannot be applied to it. Instead, we computed the free-energy change upon mutation for all 20 identities at each of the eleven strained positions, and mutations that exhibited ΔΔ*G* < +0.5 R.e.u. were selected for full combinatorial enumeration and relaxation according to the FuncLib workflow. The lowest-energy 20 designs were visually inspected and four were chosen for experimental characterization. All designs harbored eight mutations compared to the original 4H.01, and at least two mutations compared to one another. In all designs, Gln84 was mutated to Leu, alleviating the desolvation penalty of the original Gln. Furthermore, Leu78, which exhibited a strained sidechain conformation, was mutated to either Ala or Ser. The computed energies improved by at least 25 R.e.u.

The designs were expressed in pET11b vector, transformed into XL-10-Gold cells, and the proteins were purified using HisTrap^Tm^ FF column following gel filtration on Superdex 16/600 75pg. All designs showed improvement of expression relative to 4H.01 of up to threefold. The designs also improved apparent thermal stability, as determined by circular dichroism, by 17-27°C (Table 1, Fig 2C, S2). Moreover, the original 4H.01 design did not exhibit a clear melting transition, whereas all of the designs did. This suggests that the original 4H.01 design was not cooperatively folded and became a clearly folded protein upon the introduction of the eight designed mutations. We next attempted to test the designs’ ability to bind their target antigen using surface plasmon resonance (SPR). The designs, however, exhibited high binding to the reference cell, precluding accurate affinity measurements and suggesting that they bind nonspecifically. Thus, the designed mutations substantially improved stability, expressibility and apparent folding cooperativity but potentially at the cost of loss in activity. Although it remains to be seen whether the FuncLib approach can be generally applied to *de novo* designed proteins, we conclude that the pSUFER strategy flags positions that are amenable to stabilizing mutations.

**Table 1.** Expressibility and stability of FuncLib variants of *de novo* designed 4H01.

|  | Amino acid position |  |  |  |  |  |  |  | Expression levels (ug/L)<sup>1</sup> | Tm (°C)<sup>2</sup> |
| --- | --- | --- | --- | --- | --- | --- | --- | --- | --- | --- |
|  | 1 | 2 | 7 | 47 | 53 | 76 | 78 | 84 |  |  |
| 4H01 | H | M | E | H | Q | G | L | Q | 54 | 41.4 |
| 4H01_mut1 | P | E | T | T | K | R | A | L | 128 | 62.4 |
| 4H01_mut3 | P | E | T | S | K | R | S | L | 146 | 58.7 |
| 4H01_mut5 | P | E | D | R | K | R | A | L | 108 | 68.4 |
| 4H01_mut8 | P | E | T | R | K | S | A | L | 78 | 66.6 |

1. Determined using nanodrop
2. Determined by thermal melt analysis using circular dichroism (CD)

For a final example of pSUFER’s abilities and limitations, we examine binders generated through modular assembly and design. Using modular assembly and design, we previously started from a high-affinity colicin endonuclease-immunity binding pair ^40^ to design a set of binders, some of which exhibited ultrahigh specificity (>100,000 fold) relative to the parental pair and other designed pairs ^24^. Our approach focused on the design of a new interfacial loop backbone in the immunity protein by grafting loop backbones from completely unrelated proteins and optimizing the sequence of the binding pair. X-ray crystallographic analysis demonstrated that an ultraspecific designed pair (des3) exhibited atomic accuracy throughout the structure and in the designed loop relative to the model, whereas a multispecific design (des4) exhibited missing density in parts of the designed loop.

We applied pSUFER to the immunity protein in the absence of its endonuclease partner. In each design, pSUFER flags a position within the designed binding loop (Fig 3A and 3B). The flagged position in des4, Gly26, cannot be redesigned as Gly26’s backbone comes into close contact with the endonuclease. In des3, by contrast, two positions in the binding loop are flagged, Glu25 and Asn27. While Glu25 can be designed to other identities which are less strained according to our models, Asn27 forms polar contacts with the endonuclease and thus may be crucial for binding. In addition, in both designs pSUFER flags several solvent-accessible positions (Fig 3B). The results on des3 and des4 demonstrate that the pSUFER analysis may in some cases indicate problems that cannot be relieved without compromising the design of function objectives.

**Figure 3.**
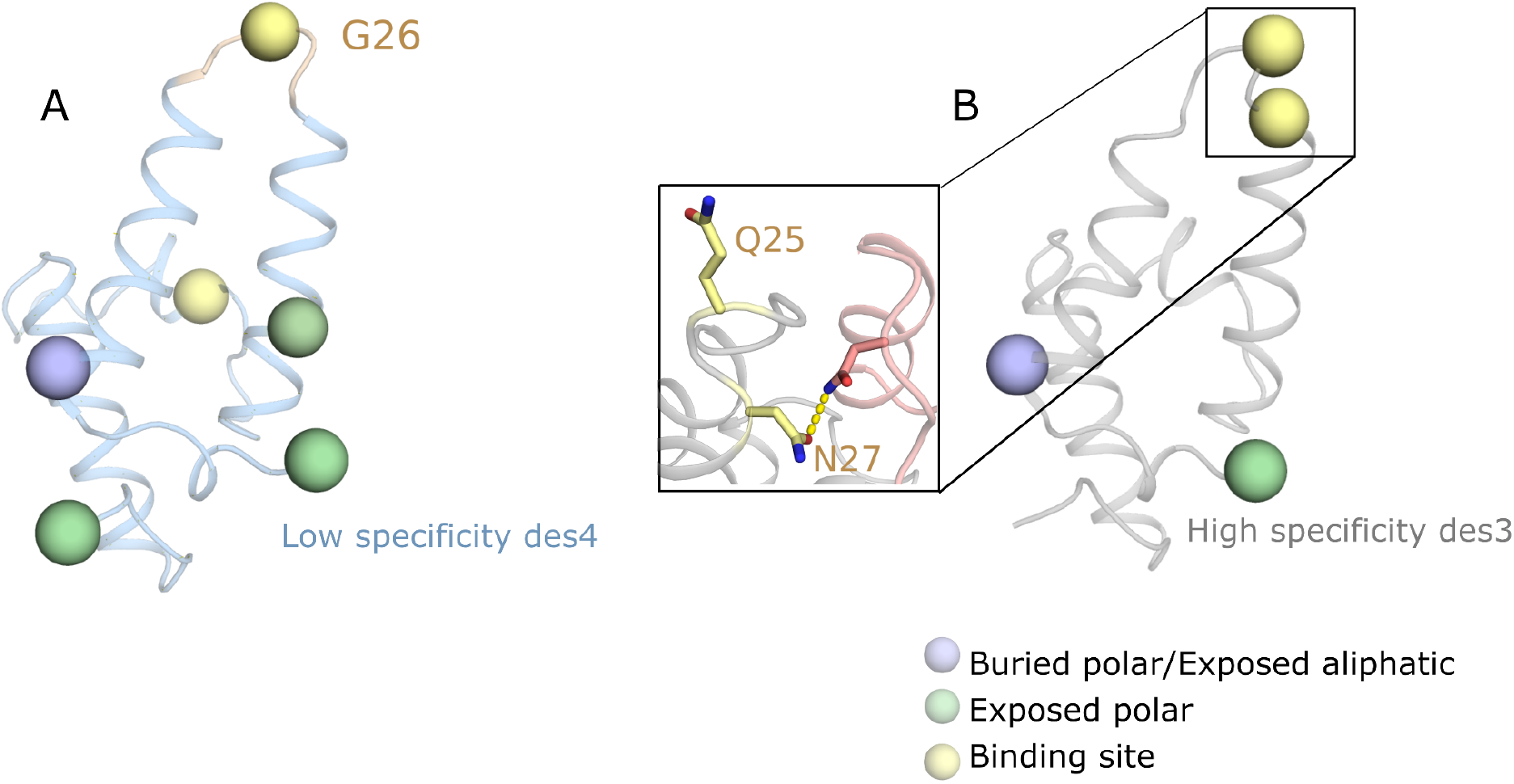
pSUFER limitations in designed binders. **(A+B)** binders generated through modular assembly and design by grafting an interfacial loop (top) from an unrelated protein and designing the two binding partners ^24^. Positions were flagged if mutational scanning suggested at least six amino acid identities with ΔΔ*G* < 0 at the position. Wheat backbone marks missing density on the design model. **(A)** The interfacial loop failed to exhibit electron density. pSUFER flags Gly26 in the designed loop. **(B)** pSUFER analysis of the accurately designed des3 flags exposed residues and two interfacial positions. Asn27 is one of the interface loop positions that forms polar contacts with the binding protein and thus may be crucial for function (inset).

**Figure 3.**
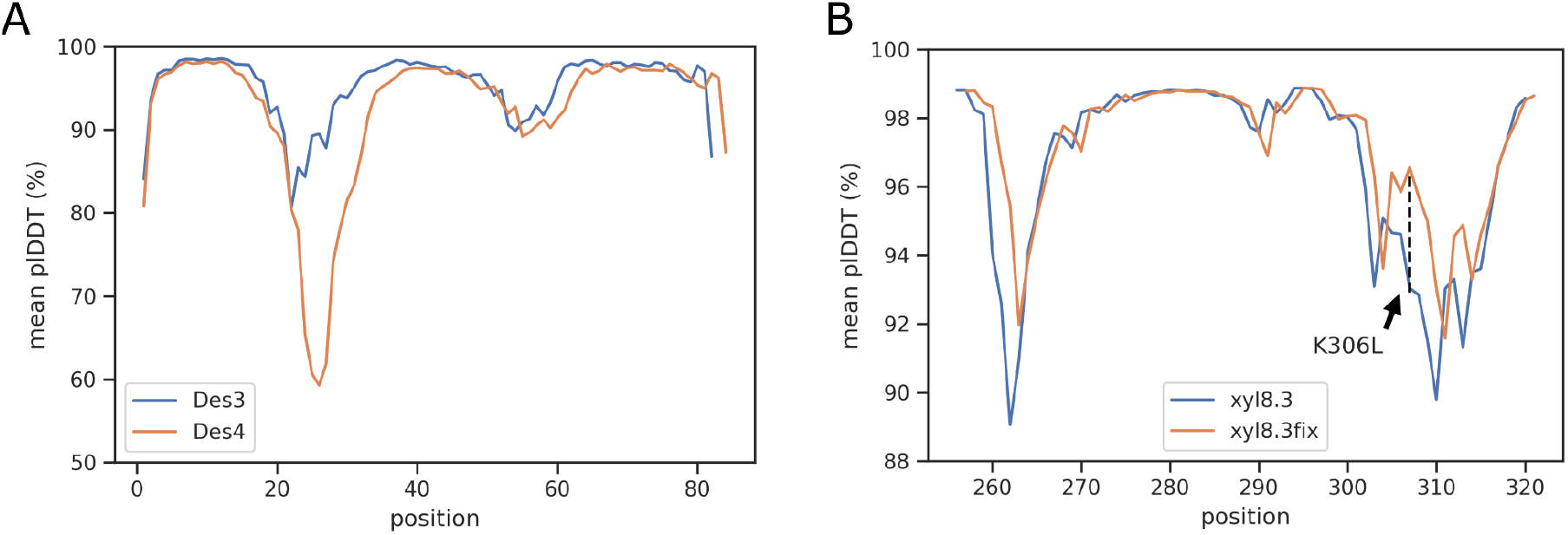
AlphaFold2 analysis of designs. **(A)** plDDT scores for designed binders des3 and des4. The plDDT scores of des4 are depressed relative to those of des3 in the region in which des4 exhibits missing density. **(B)** Comparison of the plDDT scores of xyl8.3 and design xyl8.3fix in the region surrounding the positions that exhibit missing density in crystallographic analysis of xyl8.3 (PDB entry: 6FHE; density is missing surrounding positions 259-265 and 301-312). In this region only a single mutation was designed in xyl8.3fix, Lys306Leu; yet, the plDDT scores of both loops improve significantly.

### Analyzing design foldability by AlphaFold2

pSUFER analyzes suboptimality based on a molecular structure or model and is thus limited to analyzing the native-state properties of the design. Foldability, however, also depends on whether the protein is likely to fold into alternative (misfolded) states, a possibility that may be assessed by *ab initio* structure predictors. Recently, methods that use deep learning have been very successful in *ab initio* structure prediction ^41–44^. The most successful of these, AlphaFold2, has shown atomic-accuracy prediction in community blind-prediction experiments and other structure-prediction challenges ^41^. Significantly for the purposes of assessing design foldability, AlphaFold2 provides a confidence score (per-residue local distance difference test; plDDT) for each position in the predicted model structure. We hypothesized that the plDDT scores and the rmsd between the AlphaFold2 predicted structure and the design model might predict design accuracy and foldability. AlphaFold2 depends on multiple-sequence alignments of homologs for accurate structure prediction so we did not analyze the *de novo* designed binders.

The AlphaFold2 analysis of the two accurately designed proteins (the GH10 xyl3.1 and the colicin immunity des3) correspond very well to the design models (rmsd < 0.7 Å). Furthermore, except in the N and C-terminal tails, the plDDT scores are high in xyl3.1 and the majority of des3 (>90%). For des3, the binding-surface loop backbone was grafted from a non-homologous protein, explaining why the plDDT scores are not as high in this region (>80%). Significantly, in the case of the colicin immunity des4, the AlphaFold2 model deviates from the design model in the region corresponding to the missing density (Fig 3A). Furthermore, the plDDT scores clearly depress around these regions in both des3 and xyl8.3 relative to the two other designs. We next analyzed the AlphaFold2 results for xyl8.3fix. The AlphaFold2 model structure recapitulated the design model with rmsd < 0.8Å, similar to xyl8.3. Remarkably, the plDDT scores in the region surrounding position 306, as well as in the loops that exhibit missing density according to the crystallographic analysis are higher and equivalent to those observed for the remainder of the protein (>92%) (Fig 3B). This result suggests that the AlphaFold2 confidence scores are sensitive even to single-point mutations in designed proteins. We concluded that the AlphaFold2 analysis can provide critical information on the likely accuracy and foldability of newly designed backbone structures, indicate regions that may misfold, and assess mutations that are designed to mitigate misfolding.

## Discussion

Recent advances in protein design methodology extend its scope to the design of large proteins that are highly mutated relative to natural ones ^25^ and to sets comprising thousands of designs ^3,14^. Nevertheless, the success rate is typically very low and some critical goals remain elusive. Thus, automated methods to assess design reliability may have a profound impact on the ability to design desired molecular activities. Assessing a design’s accuracy and foldability, however, remains challenging. As a general rule, design studies, particularly ones devoted to backbone design, often reveal significant deviations between experimental structures and design conceptions. When the designs are small (<100 amino acids), they may be subjected to atomistic forward-folding *ab initio* simulations to verify that the sequences favor the designed conformation over others ^1–3,45^. In large proteins, however, accurate forward-folding simulations were until recently impossible. Our results demonstrate that the deep-learning based method AlphaFold2 can shed light on design foldability even in large proteins that exhibit more than 100 mutations from any natural protein.

Furthermore, the pSUFER energy-based method can pinpoint specific positions that may be poorly designed. The functional consequences of poor design choices are strikingly demonstrated in our study: by redesigning just four positions in xyl8.3 that were flagged by pSUFER (out of 350 positions), and eight positions in 4H01 the enzyme’s catalytic efficiency rises by three orders of magnitude, and the expressibility and stability of 4H01 improves by threefold and 27 °C, respectively. These flagged positions are located outside the active or binding sites, demonstrating the significance of accurate and strain-free design throughout the protein. These observations, together with the improvement in the AlphaFold2 confidence scores for the enzyme design variant, implicate foldability or reduced active-site strain as the cause of improvement in activity. It is also striking that a handful of poor design choices in a large protein led almost to complete dysfunction. We note, however, that the protein’s ability to fold into the desired conformation also depends on other factors, such as the kinetic accessibility of the native state and the stability of folding intermediates^46^. These kinetic factors are not assessed by pSUFER, and it is unlikely that they can be deduced from current deep-learning-based *ab initio* structure predictors. Nevertheless, the strategy we described may free protein designers to introduce more radical changes than previously and increase the success of backbone design in large enzymes and binders.

## Acknowledgements

We thank Olga Khersonsky for help in xylanase activity screens and Ravit Netzer for advice in analyzing the binders. Research in the Fleishman lab was supported by a Consolidator Award from the European Research Council (815379), an Individual Grant from the Israel Science Foundation (1844/19), the Dr. Barry Sherman Institute for Medicinal Chemistry, and a charitable donation in memory of Sam Switzer. R.L-S is supported by a fellowship from the Arianne de Rothschild Women Doctoral Program. The Correia lab was supported by a Starting Grant from the European Research Council (716058) and a project grant from the Swiss National Science Foundation (310030_163139).

## Materials and methods

### pSUFER algorithm

RosettaScripts ^47^ and commandlines are available through github https://github.com/Fleishman-Lab/pSUFER. For all Rosetta calculations the ref15 energy function is used for scoring ^48^. We start the procedure with four iterations of refinement of the input structure comprising sidechain packing and harmonically constrained whole-protein minimization. Next, computational mutation scanning is performed using the FilterScan mover in Rosetta ^10^: for each position all 20 amino acids are modeled against the refined structure, and sidechains within 8 Å are repacked including constrained whole-protein minimization in order to accommodate the mutation. The energy difference between the refined structure and the single-point mutant is calculated. Positions that exhibit several mutations with ΔΔ*G* ≤ 0 R.e.u. are flagged as suboptimal with the mutation threshold in this work set to four for the *de novo* designed binders and six for all the other designs. Thresholds can be set by the user. pSUFER can be accessed through a web server at https://pSUFER.weizmann.ac.il. In this case, the user inputs a PDB-formatted coordinate file and the energy difference threshold for which a mutation is defined as favorable. The output consists of a folder containing 1) a PyMOL session file in which suboptimal positions are marked in yellow sticks. The session includes several models in which suboptimal positions are marked according to different cutoffs for the number of amino acid identities that exhibit ΔΔ*G* smaller than the set energy threshold. For instance, at threshold 4, all positions that exhibit 4 or more favorable identities relative to the starting structure are marked. 2) a table summarizing the suboptimal positions according to the cutoffs. 3) Tables indicating all of the favorable identities at each position (Rosetta resfile). 4) the Rosetta-refined structure.

### FuncLib design calculations

FuncLib on xyl8.3 was performed on the pSUFER flagged positions as described in ^12^ with the following thresholds PSSM ≥ −2 and ΔΔ*G* ≤ 1 R.e.u. in all flagged positions outside the active site. The 4H01 binder is the outcome of *de novo* design and lacks a multiple-sequence alignment from which a PSSM can be computed. We therefore extended the computational mutation scan to probe all 20 amino acid identities at any position >5 Å from the binding site, and identities that exhibited ΔΔ*G* ≤ 0.5 R.e.u were chosen for combinatorial enumeration using FuncLib. All combinations of mutations were scored using Rosetta. For xyl8.3, within the top 20 designs ranked by system energy, the designs converged on the same solution for two of the positions and we chose the design that exhibited the most polar solvent facing residues (Thr1Asn, Asn4Glu) for experimental characterization.

### AlphaFold2 *ab initio* structure prediction calculations

All AlphaFold2 ^41^ calculations were implemented by adapting and locally running the code written by ColabFold ^49^. All runs were performed using the five model parameters presented in CASP14, without templates, with Amber relaxation and using three recycle rounds only. Multiple-sequence alignments were generated through the MMseqs2 API server ^50–52^.

### Materials

4-nitrophenyl β-xylobioside (OPNPX2) was purchased from Megazyme.

### Cloning, protein expression and purification of xyl8.3

All experimental procedures were performed as described in ^25^. Briefly, the xyl8.3fix design was ordered as a synthetic gene fragment from Twist Bioscience and cloned into pETMBPH vector which contains N-terminal 6-His-tag and MBP. EcoRI and PstI sites were used. The xyl8.3 and xyl8.3fix designs were transformed into BL21 DE3 cells and the DNA was extracted and validated by Sanger sequencing.

50ml of 2YT with 50 μg ml^-1^ kanamycin was inoculated with 500 μl overnight culture and grown at 37 °C to OD of 0.4-0.8. Overexpression was induced by 0.2mM IPTG and the cultures were grown for ∼20h at 20°C. Bacteria were pelleted by centrifugation and frozen for at least 20 min before purification.

Pellets were resuspended in lysis buffer (50 mM Tris-Cl pH 6.5, 150 mM NaCl, benzonase and 0.1 mg ml^-1^ lysozyme) and lysed by sonication. The supernatant was loaded on a column packed with amylose resin (NEB), washed twice with 50 mM Tris pH 6.5 and 150 mM NaCl, and eluted with wash buffer containing 10 mM maltose. Protein purity was evaluated by SDS-PAGE gel and protein concentration was estimated by OD_280_. In cases where purity was not satisfactory, the elution was loaded on an Ni-NTA, washed and eluted (50 mM Tris-Cl pH 6.5, 150 mM NaCl, 20 mM Imidazole and 250 mM Imidazole respectively). The proteins were then dialyzed against 50 mM Tris-Cl pH 6.5, 150 mM NaCl buffer.

### Kinetic measurements of xyl8.3

Kinetic measurements were performed with purified proteins (fused to MBP) in 96-well plates (optical length –0.5 cm) by monitoring the absorbance of the leaving group of O-PNPX2 at 405 nm (activity buffer 50 mM Tris pH 6.5 and 150, 25 °C). No background hydrolysis was observed with O-PNPX2. Final protein concentrations in the reaction varied between 2.56 μM and 6.23 μM. The data were fitted to the linear regime of the Michaelis-Menten model (v_0_= [S]_0_[E]_0_k_cat_/K_M_) and kcat/KM values were deduced from the slope. The reported values represent the means ± S.D. of at least two independent measurements.

### Apparent melting-temperature measurements of enzyme designs

Tm measurements were done using nanoDSF experiments ^53^ performed on Prometheus NT.Plex instrument (NanoTemper Technologies). Melting temperatures were between 20°C-85°C with 1.0°C/min slope. For xyl8.3 and xyl8.3fix the MBP tag was removed by TEV cleavage. For xyl8.3 and xyl8.3fix the buffer was 50 mM Tris-Cl pH 6.5, 150 mM NaCl. For *de novo* enzyme designs the buffer was 100mM Tris-Cl pH 7.25, 200 mM NaCl. Protein concentrations varied between 2.8-6.0 mg/ml for all proteins. Fluorescence intensity was adjusted to suit all samples per experiment.

### Cloning, protein expression and purification of 4H01 series

The 4H01 designs were ordered as a synthetic gene fragment from Twist Bioscience with addition of a C-terminal 6-His-Tag and cloned into a pET11b vector using NdeI and BlpI restriction sites. The designs were transformed into XL-10-Gold cells and the DNA was extracted and validated by Sanger sequencing. The validated DNA sequences were transformed into BL21 DE3 cells and put in 20 ml of LB medium with 100 μg ml^-1^ Ampicillin overnight at 37 ºC as starting cultures. The next day, 500 ml of Auto-Induction medium with 100 μg ml^-1^ Ampicillin was inoculated with 10ml overnight culture and grown at 37 °C to OD of 0.6 then the cultures were grown for ∼20 h at 20 °C. Bacteria were pelleted by centrifugation and resuspended in lysis buffer (100 mM Tris-Cl pH 7.5, 500 mM NaCl, 5% Glycerol, 1mM Phenylmethanesulfonyl fluoride, 1 mg ml^-1^ lysozyme and 1:20 of CelLytic^Tm^ B Cell Lysis Reagent). The resuspentions were put at room temperature on a shaker at 40 rpm for 2 hours and then centrifuged at 48,300g for 20 minutes. We filtered the supernatant with a 0.2 μm filter and loaded the mixture on a 1ml HisTrap^Tm^ FF column using an AKTApure system and a predefined method regarding Cytiva’s recommendations with that column. We used 50 mM Tris-HCl pH 7.5, 500 mM NaCl, 10 mM Imidazole as wash buffer and processed the elution with 50 mM Tris-HCl pH 7.5, 500 mM NaCl, 500 mM Imidazole. We collected the main fraction released through the elution step and injected it on a Gel Filtration column Superdex 16/600 75pg filled with PBS. The peaks corresponding to the size of the design were collected and concentrated till a concentration of approximately 1mg ml^-1^ for further analysis. Protein concentrations were determined by nanodrop.

### Apparent melting-temperature measurements of 4H01 series

Tm measurements were done using Chirascan^Tm^ V100 from appliedPhotophysics. Melting temperatures were between 20-90 °C with measurements every 2 ºC.

## Supporting Information

### Supplementary material

**Supplementary Figure 1.**
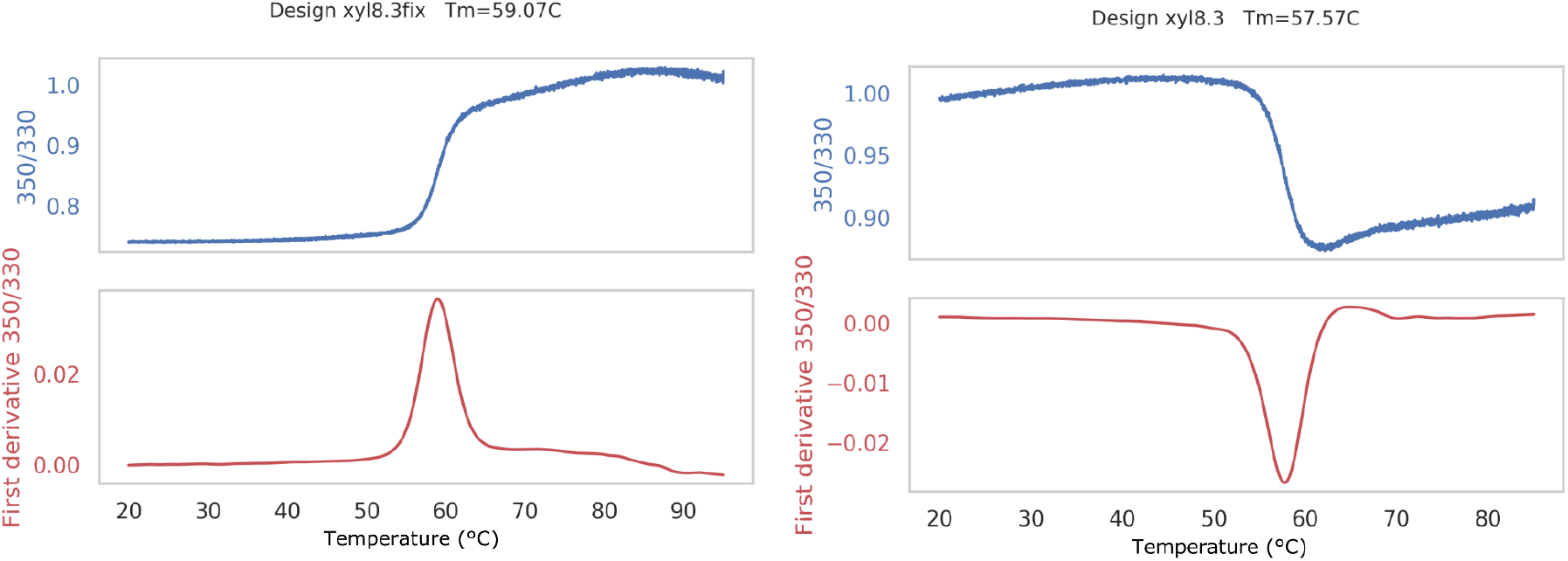
Denaturation curves for xyl8.3 and xyl8.3fix (blue - ratio, red - derivative) using nanoDSF.

**Supplementary Figure 2.**
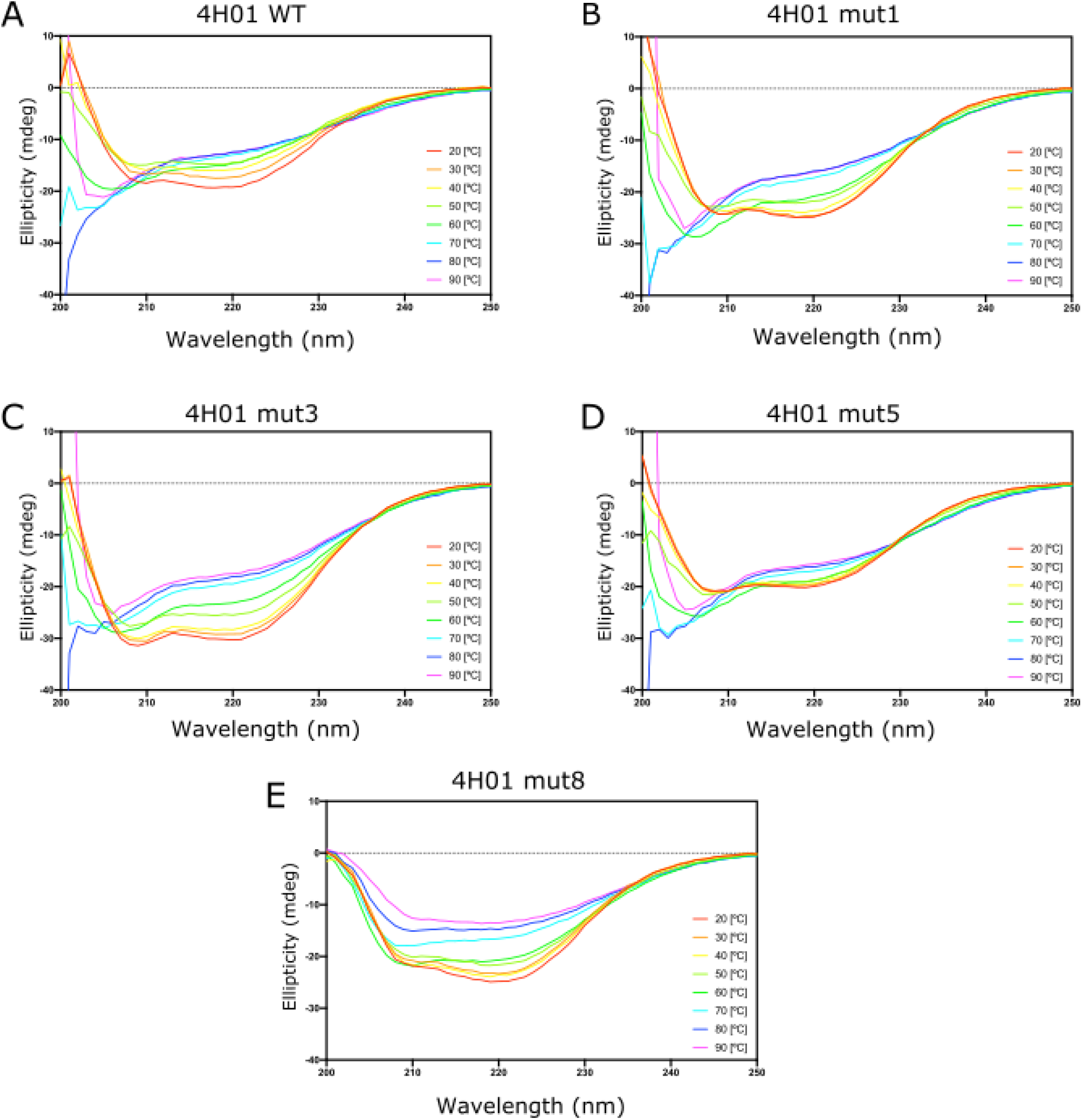
CD spectra of the 4H01 binder and four FuncLib mutants (A-E) at temperatures ranging 20-90 °C.

**S3 Fig**. Temperature melts of 4H01 variants monitored by CD at 220 nm

